# Single-cell transcriptomics identifies CD44 as a new marker and regulator of haematopoietic stem cells development

**DOI:** 10.1101/338178

**Authors:** Morgan Oatley, Özge Vargel Bölükbasi, Valentine Svensson, Maya Shvartsman, Kerstin Ganter, Katharina Zirngibl, Polina V. Pavlovich, Vladislava Milchevskaya, Vladimira Foteva, Kedar N. Natarajan, Bianka Baying, Vladimir Benes, Kiran R. Patil, Sarah A. Teichmann, Christophe Lancrin

## Abstract

The endothelial to haematopoietic transition (EHT) is the process whereby haemogenic endothelium differentiates into haematopoietic stem and progenitor cells (HSPCs). The intermediary steps of this process are unclear, in particular the identity of endothelial cells that give rise to HSPCs is unknown. Using single-cell transcriptome analysis and antibody screening we identified CD44 as a new marker of EHT enabling us to isolate robustly the different stages of EHT in the aorta gonad mesonephros (AGM) region. This allowed us to provide a very detailed phenotypical and transcriptional profile for haemogenic endothelial cells, characterising them with high expression of genes related to Notch signalling, TGFbeta/BMP antagonists (Smad6, Smad7 and Bmper) and a downregulation of genes related to glycolysis and the TCA cycle. Moreover, we demonstrated that by inhibiting the interaction between CD44 and its ligand hyaluronan we could block EHT, identifying a new regulator of HSPC development.

## Introduction

Understanding the developmental origin of haematopoietic stem and progenitor cells (HSPCs) is of critical importance to efforts to produce blood and blood products *in vitro* for medical applications. Haematopoietic stem and progenitor cells (HSPCs) originate from endothelial cells in the aorta gonad mesonephros (AGM) of midgestation embryos ^1,2^. This process known as the endothelial to haematopoietic transition (EHT) requires drastic morphological changes that have been directly visualised through time-lapse imaging studies both in vitro and in vivo ^3-6^. EHT is a highly conserved process that has been studied across vertebrate models from xenopus and zebrafish to mice ^7^. Importantly, the human definitive blood system also has an endothelial origin ^8^.

The best tools so far to detect endothelial cells with haemogenic capabilities rely on using fluorescent reporters under the control of Runx1^9^ or Gfi1 ^10^ regulatory elements, two keys transcription factors in the process of EHT. Cells expressing these transcription factors already co-express blood and endothelial genes. However, we still do not know the nature of endothelial cells which will acquire the expression of these transcription factors. We still do not know if any endothelial cells in the AGM can initiate the haematopoietic program or if a certain type of endothelial cells is primed to undergo the EHT.

Despite our lack of characterisation of the definitive precursor to HSPC development, haemogenic endothelium (HE), recent advances have been made in terms of *in vitro* HSPC generation via an endothelial intermediate. Through the use of transcription factor cocktails both human pluripotent stem cells via a HE stage and adult mouse endothelial cells have been successfully reprogrammed into multi-potent, definitive haematopoietic stem cells (HSCs) ^11,12^. However, the use of endothelial populations in the process emphasises the importance of an improved understanding of HE.

The early haematopoietic hierarchy has been described as a three-step process based on phenotypic characteristics. Specifically, Pro-HSC, Pre-HSC type I and type II populations have been defined based on their expression of the cell surface markers CD41, CD43 and CD45, as well as the time taken mature into definitive HSCs in OP9 co-culture ^13^. Recently, an in depth transcriptional investigation was performed on the type I and type II Pre-HSC populations at day 11 of mouse embryonic development, however, the earlier stages of EHT remain largely uncharacterised ^14^. Indeed, gaining a solid understanding of HE and the initial steps that endothelial cells must take to become HSPCs has proved difficult in the absence of a robust marker.

Through antibody screening and single-cell RNA sequencing (sc-RNA-seq) we identified CD44 as a novel marker of EHT, enabling the isolation of key cellular stages of blood cell formation in the embryonic vasculature. CD44 is a cell surface receptor principally involved in the binding of the extracellular matrix molecule hyaluronan ^15^. Its cell surface expression has been used to identify cancer stem cell populations and has been strongly linked to the metastatic potential of many cancers ^16-20^. While previous research has revealed the importance of CD44 in HSPCs migration to the bone marrow, the role of the receptor in early embryonic haematopoiesis has not been characterised ^21^. Using CD44 expression in conjunction with VE-Cadherin (VE-Cad) and Kit we could clearly differentiate between vascular endothelium, HE, Pre-HSPC-I and Pre-HSPC-II more accurately than using the combination of VE-Cad, CD41, CD43 and CD45 markers. This has allowed us to perform extensive transcriptional profiling making it possible to characterise the very earliest changes in haematopoietic differentiation from endothelial cells. Moreover, by disrupting the interaction of CD44 and its ligand, we could inhibit EHT, demonstrating an unexpected role for CD44 in the emergence of HSPCs.

## Results

### Antibody screening and single-cell RNAseq identifies CD44 as a potential marker of early haematopoietic fate

It is well established that HSPCs have an endothelial origin within the embryo ^4-6,22^. In order to better characterise the transition between endothelial and haematopoietic identity, we performed both an *in vitro* antibody screen and an *in vivo* single-cell RNAseq experiment to identify new markers allowing the identification of subpopulations within VE-Cad^+^ cells (Supplementary Fig. S1). Antibodies against 176 cell surface markers ^23^ were tested against the VE-Cad^+^ population generated from our *in vitro* ESC differentiation system into blood cells which recapitulates faithfully the early blood development process ^24^. Forty-two of these markers were expressed on VE-Cad^+^ cells (Supplementary Table S1). We looked for bimodal expression to separate distinct endothelial populations and identified a short-list of sixteen candidates to test *in vivo* including CD41 and CD117 (also known as Kit, a marker of HSPCs) already known to split VE-Cad^+^ cells ^13^. Eight of these markers (CD44, CD51, CD55, CD61, CD93, Icam1, Madcam1 and Sca1) were found to split the VE-Cad^+^ endothelial population of the AGM in two (Fig. 1a). In parallel, VE-Cad^+^ cells were isolated from the AGM region of E10.5 embryos and their transcriptional profiles analysed sc-RNA-seq (Fig. 1b-d). Clustering analysis identified a population with both haematopoietic and endothelial gene expression, distinct from the other endothelial population (Fig. 1d). Bioinformatics analysis showed that *Cd44* is one of the best marker genes for this population of transitioning cells co-expressing endothelial and haematopoietic genes (Fig. 1c). The expression of *Cd44* was also positively correlated with other known haematopoietic markers such as *Runx1, Gfi1* and *Adgrg1 (Gpr56)* (Fig. 1d).

**Figure 1:**
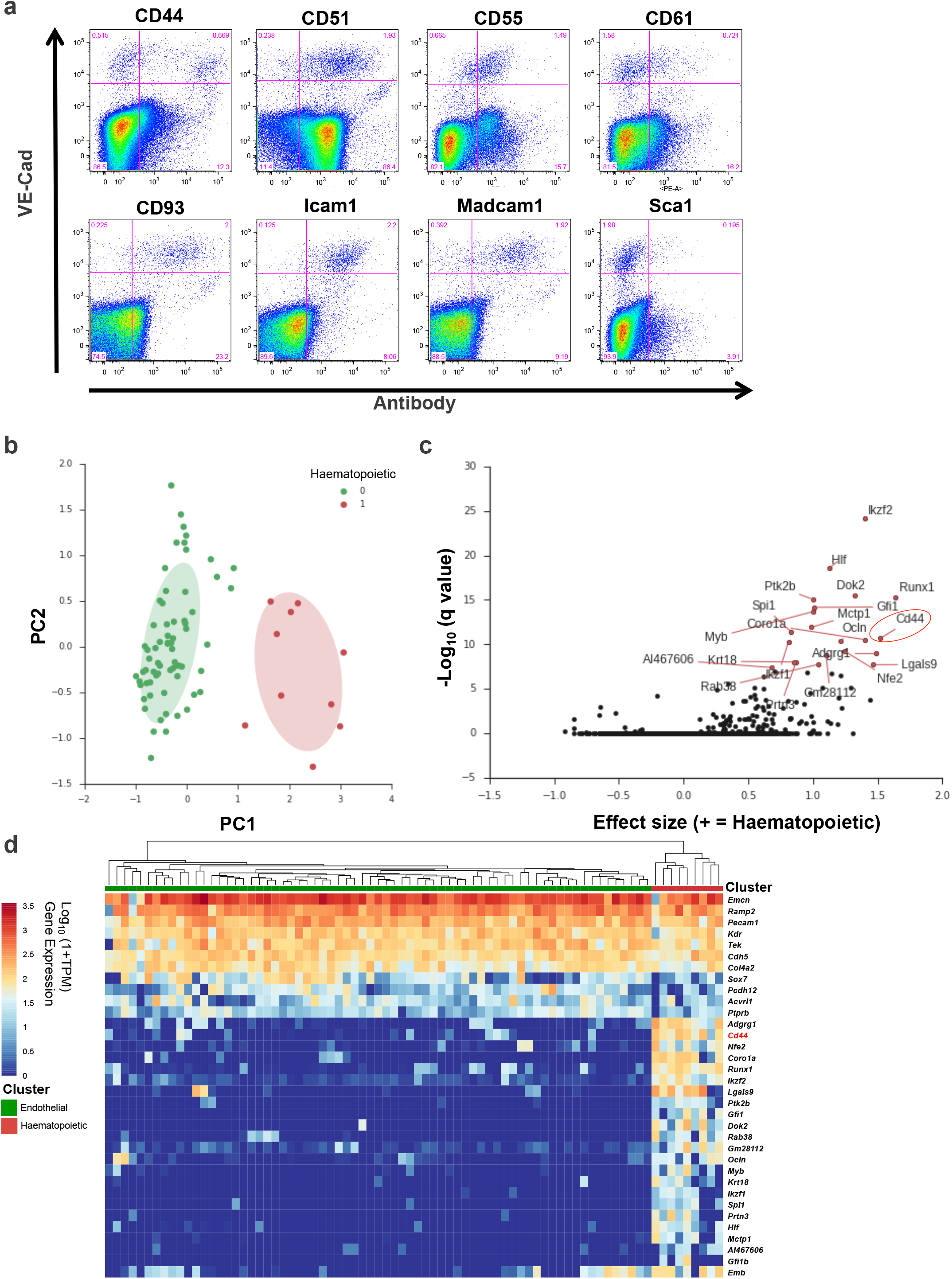
Search for a new marker to dissect the endothelial to hematopoietic transition. **(a)** FACS plot of cells isolated from AGM region at E11, stained with VE-Cad and indicated cell surface markers selected from the antibody screen. **(b)** Principal Component Analysis of the single-cell RNA-seq data. Cells expressing hematopoietic genes are marked in red while the other cells are marked in green. **(c)** Volcano plot showing a selection of marker genes specific of the groups of cells expressing haematopoietic genes. The *Cd44* gene is highlighted with a red circle. **(d)** Heatmap displaying the expression of a selection of genes in endothelial and haematopoietic clusters. *Cd44* is highlighted in red. See also Supplementary Figure S1 and File S1.

Given the association of *Cd44* with endothelial cells undergoing EHT at both the protein and mRNA level we decided to further investigate its role in embryonic haematopoiesis.

### CD44 marks transitioning cells with differing morphology

To validate our screening results and investigate the identity of CD44^+^ cells, we performed immunofluorescence and more detailed flow cytometry analysis on the AGM region of mouse embryos at embryonic day 9.5 (E9.5) and 10.5 (E10.5) (Fig. 2). Immunofluorescence of cross-sections of mouse AGMs revealed that CD44 marked cells that were part of the vascular wall and cells that were incorporated in haematopoietic clusters (Fig 2a). Flow cytometry revealed that CD44 expression significantly increased in the VE-cad^+^ endothelium of the AGM between E9.5 and E10.5 when cells are undergoing EHT (Fig. 2b-c). Furthermore, by staining with an antibody against Kit (a marker of intra-aortic haematopoietic clusters), we found that the majority of cells with lower levels of CD44 expressed little or no Kit (Fig 2d).

**Figure 2:**
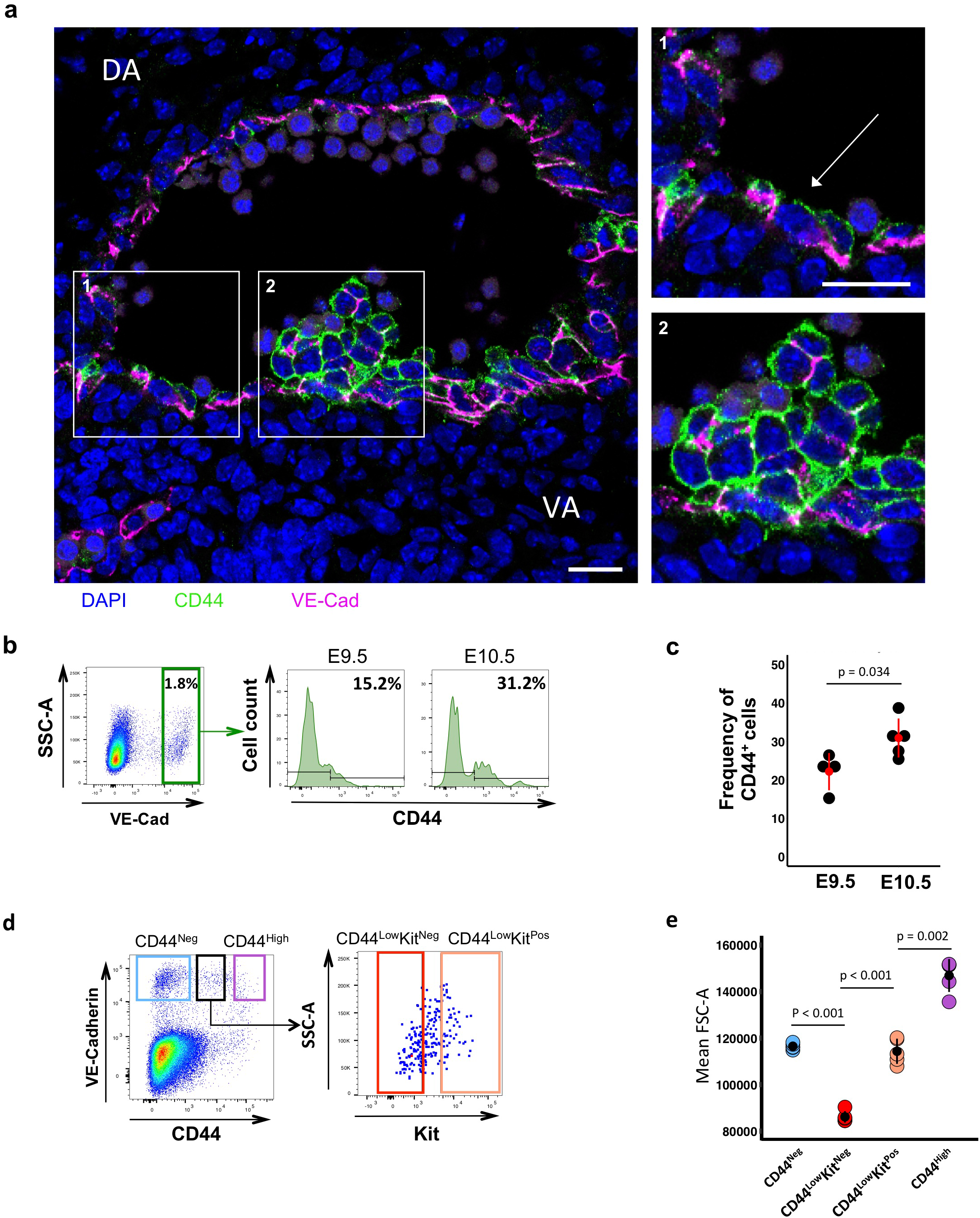
CD44 splits the AGM VE-Cadherin^+^ cells in different populations with various morphologies. **(a)** Immunofluorescence of VE-Cad (magenta) and CD44 (green) expression in a cross section of the AGM region of a wild-type embryo at E10 (32 somite pairs). Images 1 and 2 show higher magnification of the areas highlighted in the main image, showing CD44 marking endothelial cells in the vascular wall and cells making up a haematopoietic cluster. **(b)** FACS plots indicating percentage of cells expressing high levels of VE-Cad from dissected AGMs of wild-type embryos. The histograms indicate the percentage of VE-Cad^High^ cells positive for CD44 at both E9.5 (28 somite pairs) and E10.5 (35 somite pairs). **(c)** Percentage of CD44^+^ cells within the VE-Cad^High^ fraction, each data point represents an independent experiment, E9.5 n = 4 and E10.5 n = 5. Significance was determined by two-tailed, independent t-test. **(d)** FACS plots of VE-Cad and CD44 expression in the AGM region of wild-type embryos at E10 (30-34 somite pairs). Expression of Kit cell surface marker is highlighted for the CD44^Low^ population. **(e)** Mean FSC-A as an indication of cell size is plotted for each population (CD44^Neg^, CD44^Low^Kit^Neg^, CD44^Low^Kit^Pos^, CD44^High^) for five independent experiments on a litter of E10 wild-type embryos. Significance was determined by two-tailed, paired t-tests.

By grouping the cells based on their cell surface expression of CD44 and Kit, we found these populations to be significantly different in terms of cell size, suggestive of cells undergoing a morphological transition (Fig. 2e). Altogether, our results indicate that CD44 marks a subset of endothelial cells and cells in the haematopoietic clusters of the AGM during the key window of HSPCs development in the mouse embryo.

### Single-cell q-RT-PCR analysis identifies three homogeneous CD44^+^ populations with an increasing haematopoietic profile

Using the Biomark HD single-cell qPCR platform, we analysed the expression of 95 genes associated with both endothelial and haematopoietic cell types ^24^. We performed extensive transcriptional profiling on the CD44^Neg^, CD44^Low^Kit^Neg^, CD44^Low^Kit^Pos^ and CD44^High^ populations identified (Fig. 2d) between E9.5 and E11.5 (342 cells in total). We found that the CD44^Low^Kit^Pos^ and CD44^High^ populations expressed numerous haematopoietic genes (Fig. 3a). The CD44^Low^Kit^Pos^ cells also expressed many endothelial genes as in our sc-RNA-seq analysis (Fig. 1d), however the CD44^High^ cells appeared to be more advanced in the EHT process and lacked endothelial gene expression (Fig. 3a). Of note, the CD44^Low^Kit^Pos^ population expressed specifically *Gfi1* and *Itgb3* (coding for CD61) as well as the heptad of transcription factors *(Gata2, Runx1, Lyl1, Erg, Fli1, Lmo2* and *Tal1*) whose simultaneous co-expression is responsible for the dual endothelial-haematopoietic identity of Pre-HSPCs (Supplementary Fig. S2) ^24^.

**Figure 3:**
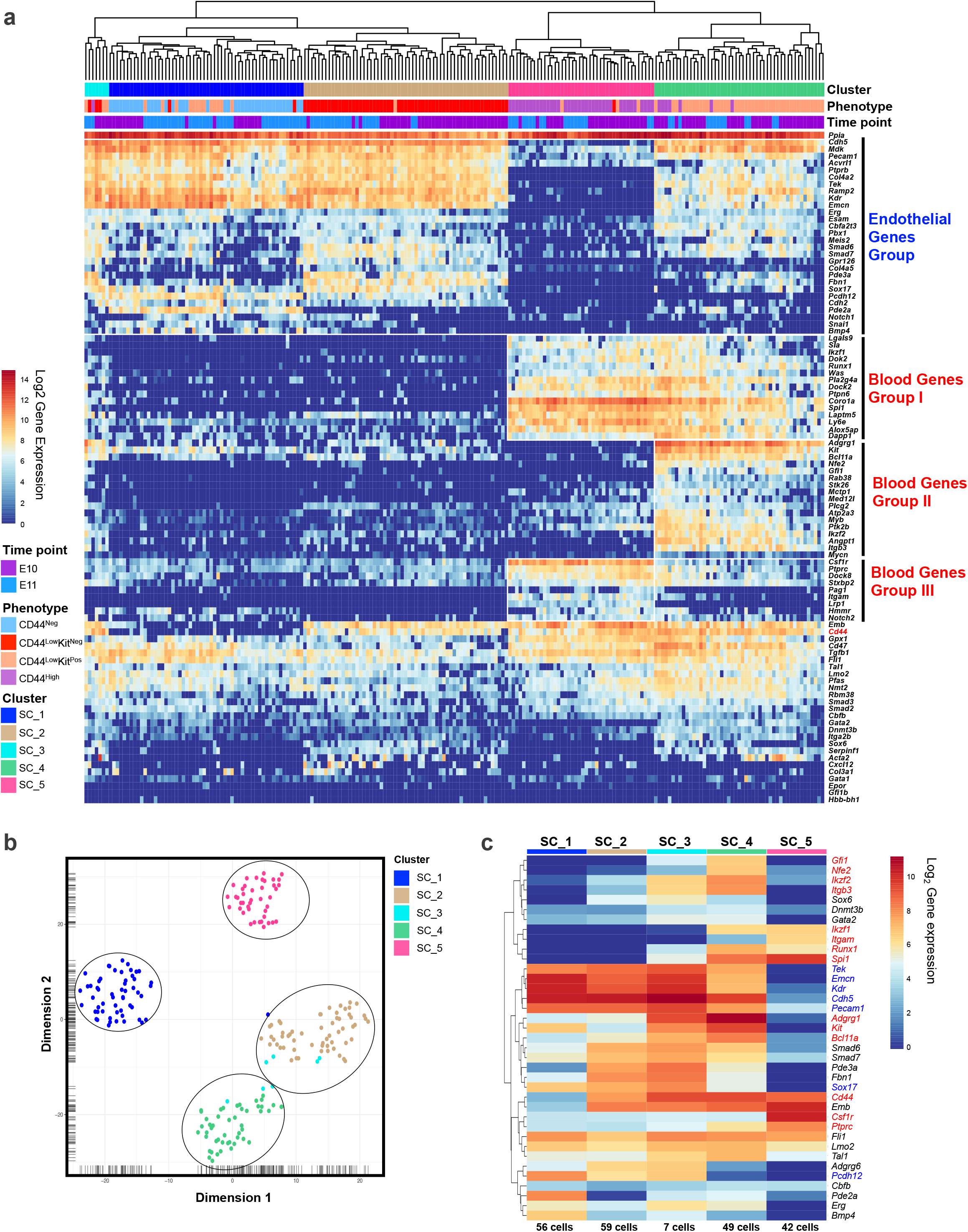
The VE-Cad^+^ subpopulations defined by CD44 are transcriptionally distinct. **(a)** Single cells from each VE-Cad^+^ populations were isolated and tested for the expression of 96 genes by single-cell q-RT-PCR. The heatmap shows the result of the hierarchical clustering analysis (cells were clustered by Euclidian distance). A selection of genes has been highlighted: endothelial genes group, blood genes groups I, II and III. **(b)** tSNE plot from single-cell q-RT-PCR data shown in (a). Five groups are indicated. **(c)** Heatmap showing average expression of endothelial (blue), haematopoietic (red) and various (black) genes in the indicated groups (genes are clustered using Pearson correlation). The number of cells for each cluster is indicated at the bottom of the panel. See also Supplementary Fig. S2 and File S3.

Conversely, the CD44^Neg^ and CD44^Low^Kit^Neg^ populations both showed specific endothelial gene signatures and lacked haematopoietic gene expression (Fig. 3a). Despite their different CD44 expression patterns, these cell populations clustered together (Supplementary Fig. S2). We repeated this experiment using a new selection of 96 genes based on our sc-RNA-seq experiment (Supplementary Table S2). With our new gene list, we were able to confirm the dual endothelial-haematopoietic and haematopoietic identities of the CD44^Low^Kit^Pos^ and CD44^High^ populations, respectively. Surprisingly, the CD44^Neg^ and CD44^Low^Kit^Neg^ populations formed two distinct clusters (Fig. 3b). In addition to *Cd44,* we found the genes *Adgrg6, Emb, Fbn1, Pde3a, Plcg2, Serpinf1, Smad6, Smad7, Sox6* and *Stxbp2* to be up-regulated in the CD44^Low^Kit^Neg^ cells compared to CD44^Neg^. In contrast, *Bmp4, Kit, Hmmr* and *Pde2a* were more expressed in CD44^Neg^ endothelial cells (Fig. 3c).

Although the four groups defined by VE-Cad, CD44 and Kit were confirmed to be distinct transcriptionally, our clustering analysis found a fifth population (SC_3) composed of cells from both CD44^Low^Kit^Neg^ and CD44^Low^Kit^Pos^ groups (Fig. 3a). Its transcriptional profile was found to be intermediary between SC_2 (CD44^Low^Kit^Neg^) and SC_4 (CD44^Low^Kit^Pos^) e.g. it expressed *Adgrg1, Runx1, Itgb3* and *Spi1* like SC_4 but was still expressing *Adgrg6* and *Pcdh12* like SC_2 (Fig. 3c).

This suggests a developmental link between the CD44^Low^ populations where CD44^Low^Kit^Neg^ cells would be the direct precursors of the CD44^Low^Kit^Pos^ population which would then go on to generate CD44^High^ cells. Interestingly, it is within this transitional SC_3 population that we saw the up regulation of *Runx1, Spi1* and *Gfi1,* which are three of the four transcription factors used to reprogram adult mouse endothelial cells into HSCs ^12^.

### VE-Cad, CD44 and Kit could segregate the earliest stages of haematopoietic development more accurately than the combination of VE-cad, CD41, CD43 and CD45 markers

To place our results in context of known AGM subpopulations, we performed transcriptional analysis of the Pro-HSC, Pre-HSC type I, and Pre-HSC type II populations defined according to the combination of VE-cad, CD41, CD43 and CD45 markers (Fig. 4a) ^13^. Following hierarchical clustering, we found three clusters: the first mostly composed of Pro-HSCs, a second being a mix of Pro-HSCs and Pre-HSCs type I and a third one composed only of Pre-HSCs type II (Supplementary Fig. S3). We then performed a clustering analysis in conjunction with the populations defined by CD44 (Fig. 4b). This revealed that the SC_5 (CD44^High^) population closely associated with the Pre-HSC type II and the SC_4 (CD44^Low^Kit^Pos^) population with Pre-HSC type I and part of the Pro-HSCs. SC_2 (CD44^Low^Kit^Neg^) clustered closely with the remaining Pro-HSCs. Finally, only five cells with Pre-HSCs type I and Pro-HSCs phenotype clustered with the SC_1 (CD44^Neg^). Ninety-seven per cent of Pro-HSCs, Pre-HSCs type I, and Pre-HSCs type II were CD44 positive (Supplementary Fig. S3 and Fig. 4c).

**Figure 4:**
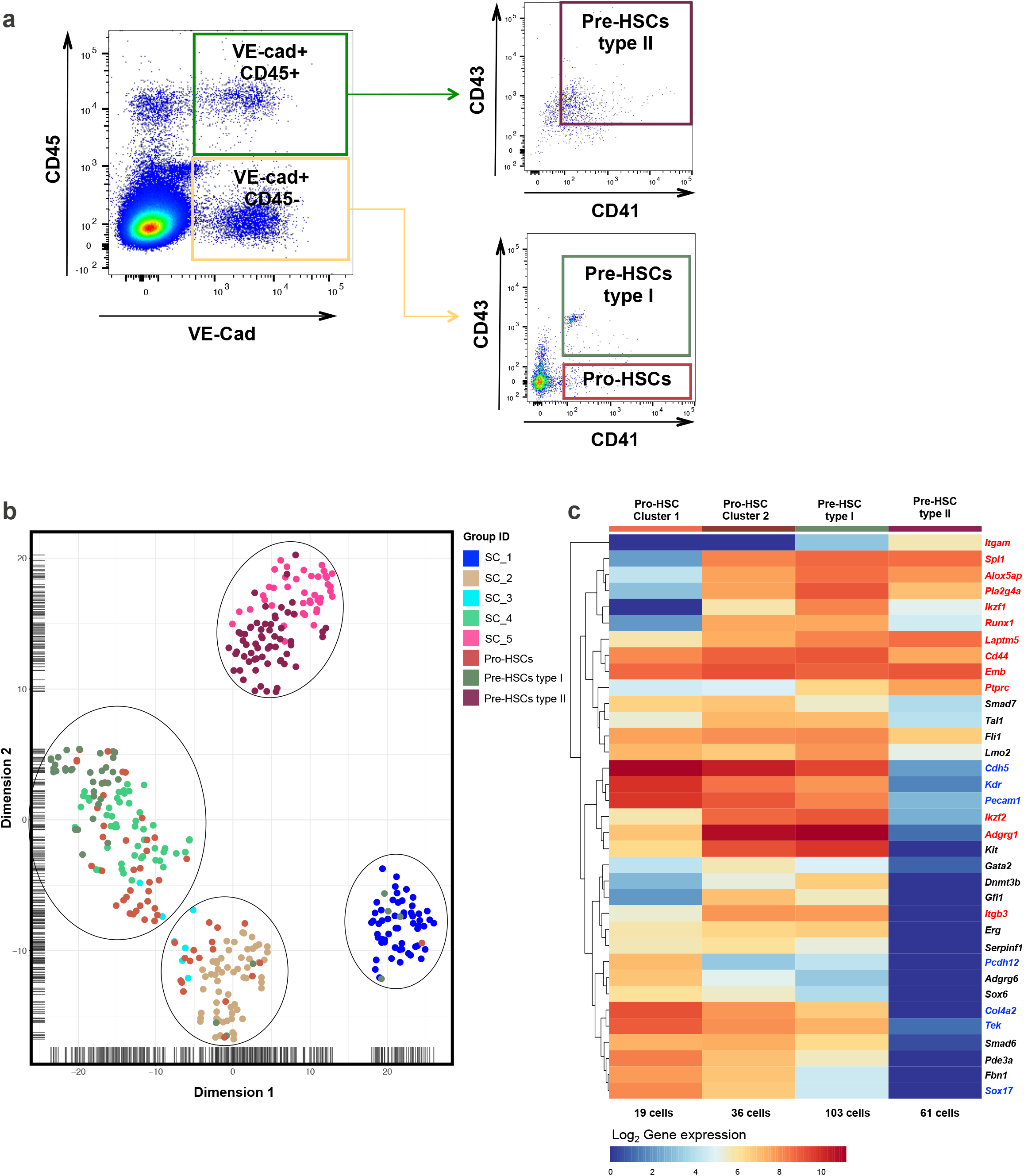
Comparison of the CD44 populations to Pro-HSC, Pre-HSC type I and type II. (**a**) FACS plots of VE-Cad, CD45, CD43 and CD41 expression in the AGM region at E10 (31-32 somite pairs). Single cells from Pro-HSCs (VE-Cad^+^ CD41^+^ CD45^-^ CD43^-^), Pre-HSCs type I (VE-Cad+ CD41+ CD45^-^ CD43+), Pre-HSCs type II (VE-Cad^+^ CD45^+^) populations were isolated. (**b**) tSNE plot from single-cell q-RT-PCR data shown in (a). (**c**) Heatmap showing average expression of endothelial (blue), haematopoietic (red) and various (black) genes in the indicated groups (genes are clustered using Pearson correlation). The number of cells for each cluster is indicated at the bottom of the panel. See also Supplementary Fig. S3 and File S4.

Overall, we have demonstrated that the phenotypes based on CD44 expression could allow us to isolate all key populations in the process of HSCs formation more accurately than the phenotypes previously described.

### Bulk RNA-seq analysis further distinguishes CD44^Neg^ and CD44^Low^Kit^Neg^ endothelial populations and identifies early changes in the differentiation process

To further compare the two endothelial populations found in the AGM, we performed RNAseq on 25-cell bulk samples from these populations across three different time points (E9.5, E10 and E11). We analysed as well the more advanced stages in EHT: CD44^Low^Kit^Pos^ at E9.5 and E10 and CD44^High^ at E11. The bulk RNA-seq approach allowed us to detect low abundant genes (such as genes encoding transcription factors) more efficiently than sc-RNA-seq and also to measure smaller changes in gene expression between populations.

The samples clustered according to their marker expression, despite the difference in developmental time, confirming our previous experiments (Fig. 5a). We identified several haematopoietic genes switched on in the CD44^Low^Kit^Neg^ population including, *Ctsc, Nfe2, Runx1* and *Ifitm1,* suggesting that these cells are subjected to the EHT process (Fig. 5b). The expression pattern of endothelial genes fits with our previous observations – more highly expressed in CD44^Neg^ and CD44^Low^Kit^Neg^, moderately expressed in CD44^Low^Kit^Pos^ and absent in CD44^High^ (Fig 5b). Moreover, we used this dataset to check the expression of genes corresponding to proteins we found in our antibody screen (Fig. 1a). *Cd93* and *Madcam1* showed an endothelial expression pattern and could be used to separate endothelial from blood cells in the AGM. In contrast, *Itgb3* marks specifically the CD44^Low^Kit^Pos^ population while Ly6e marks both CD44^Low^Kit^Pos^ and CD44^High^ populations as shown previously in Supplementary Fig. S2 and Fig. 3.

**Figure 5:**
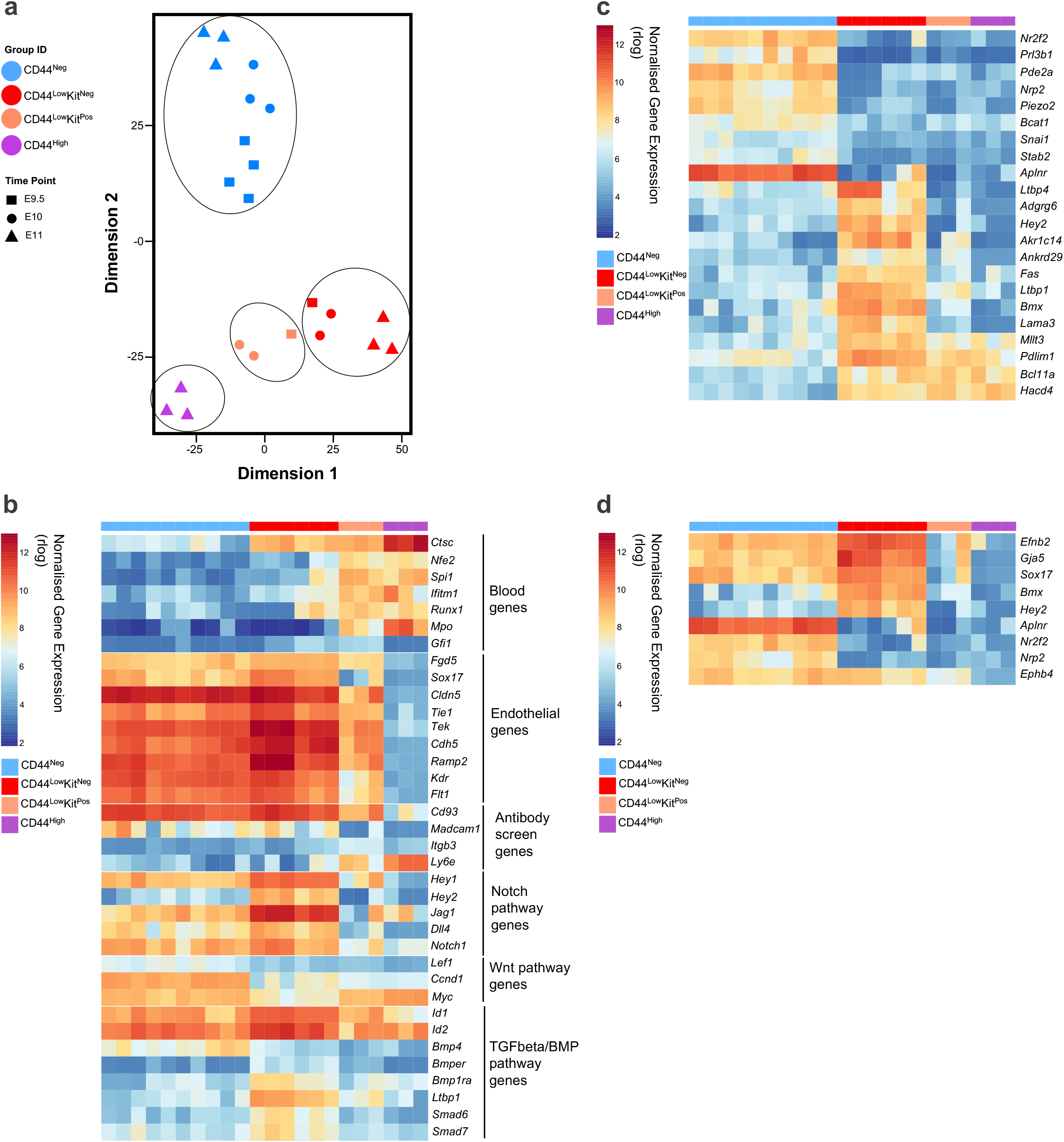
Bulk RNA sequencing further distinguishes CD44^Neg^ and CD44^Low^Kit^Neg^ endothelial populations and identifies early changes in the differentiation process. (**a**) tSNE plot from 25-cell bulk RNA sequencing generated from E9.5, E10 and E11 AGM. Four groups are indicated. (**b**) Heatmap of gene expression highlighting a selection of genes. (**c**) Heatmap of gene expression highlighting the most differentially expressed genes between CD44^Neg^ and CD44^Low^Kit^Neg^ endothelial populations (p-value < 0.01). (**d**) Heatmap of gene expression highlighting arterial *(Efbn2, Gja5, Sox17, Bmx* and *Hey2*) and venous *(Aplnr, Nr2f2, Nrp2* and *Ephb4*) coding genes in CD44^Low^Kit^Neg^ and CD44^Neg^ populations. See also Supplementary File S5 and Fig. S4.

Interestingly, this transcriptome analysis showed strong differences between the two endothelial populations of the AGM. We found 1605 genes differentially expressed between these two populations (p-value < 0.01, Wald test). Among them, several genes from the Notch pathway *(Hey2, Jag1, Dll4, Hey1* and *Notch1)* were significantly more expressed in the CD44^Low^Kit^Neg^ population compared to CD44^Neg^. Similarly, antagonists of the TGFbeta/BMP pathway, including *Smad6, Smad7 and Bmper,* were up-regulated in CD44^Low^Kit^Neg^ cells compared to CD44^Neg^ ones. In contrast, target genes of the Wnt pathway *(Lef1, Ccnd1* and *Myc)* were more highly expressed in CD44^Neg^ compared to CD44^Low^Kit^Neg^. From this analysis we could also identify specific markers for each of the endothelial populations (Fig. 5c). *Nr2f2, Pde2a, Aplnr* and *Kcne3* marked specifically CD44^Neg^ cells while *Adgrg6, Hey2, Akr1c14, Fas* and *Ltbp1* strongly marked the CD44^Low^Kit^Neg^ population. Sc-RNA-seq done by another group investigated the different stages of EHT in the AGM ^14^, but they did not identify the CD44^Low^Kit^Neg^ population. However, the gene expression pattern they obtained for the three other populations was very similar to the one we found with our bulk RNA analysis (Supplementary Fig. S4).

Our data supports the hypothesis that the CD44^Neg^ cells and the CD44^Low^Kit^Neg^ cells belong to two distinct endothelial populations. The CD44^Neg^ population expresses venous *(Aplnr, Nr2f2* and *Nrp2)* and arterial *(Sox17, Bmx* and *Efnb2)* markers while the CD44^Low^Kit^Neg^ has a clear arterial signature with stronger expression of *Bmx*, *Jag1* and *Hey2*. Moreover, there are distinct transcriptional links between the two CD44^Low^ populations and the initiation of several blood markers already at the CD44^Low^Kit^Neg^ stage suggests that this population is in fact the endothelial precursor of CD44^Low^Kit^Pos^ and CD44^High^ cells, and hence of haematopoietic development.

### The CD44^Neg^ and CD44^Low^Kit^Neg^ endothelial populations feature distinct metabolism and autophagy signatures

Remarkably, a large number of the differentially expressed genes between the two endothelial populations belonged to metabolic processes (395 out of 1605 genes; 1.32-fold enrichment, p-value <0.05, Fisher’s exact test). We therefore identified specific metabolic pathways distinguishing the two populations (Fig. 6a and Supplementary Table S3). Notably, the CD44^Low^Kit^Neg^ population showed a pronounced down regulation of genes coding for enzymes involved in glycolysis, TCA cycle and respiration, suggesting reduced ATP generation. Furthermore, amino acid and nucleotide biosynthesis genes were also down regulated. Altogether it suggests that the CD44^Low^Kit^Neg^ is a non-proliferative, metabolically rather quiescent state which is in line with the smaller size of this population compared to the CD44^Neg^ (Fig. 2e). Given that endothelial cells are known to obtain most of their energy from glycolysis ^25^, this change in metabolic status supports a loss of endothelial identity. These cells also show a marked increase in the expression of pathways leading to lipids with regulatory function: glycerolipids, glycerophospholipids, phosphatidylinositol and sphingolipids (Fig. 6a).

**Figure 6:**
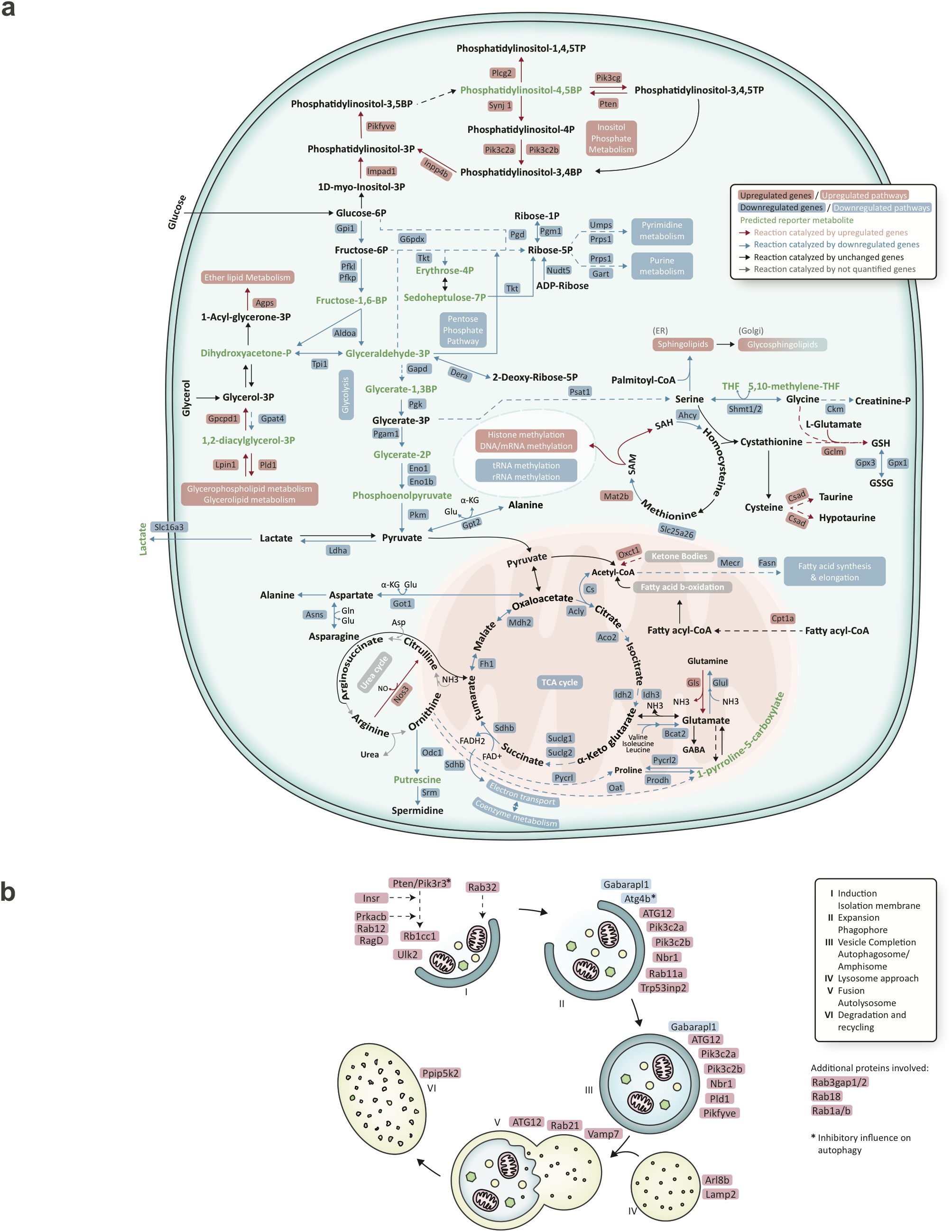
CD44^Low^Kit^Neg^ endothelial cells feature altered expression of genes involved in metabolism and autophagy. (**a**) Overview of key metabolic nodes and pathways enriched in differentially expressed genes when comparing the CD44^Low^Kit^Neg^ and CD44^Neg^ endothelial populations. These were selected based on reporter metabolite analysis. Pathway boxes summarize multiple genes / reporter metabolites (Supplementary Table S3). The mentioned upregulated and downregulated genes refer to the expression in the CD44^Low^Kit^Neg^ population compared to CD44^Neg^. (**b**) Schematic representation of the autophagy process marking differentially expressed genes. Included are genes coding for structural as well as regulatory aspects of autophagy. The colour code is the same as in (a).

Interestingly, we observe that several genes involved in autophagy, a process known to be regulated by phosphatidylinositol and sphingolipids ^26-29^, are also highly upregulated in the CD44^Low^Kit^Neg^ population (Fig. 6b). Concordantly, two key processes accompanying autophagy, ubiquitylation and proteolysis, are also upregulated. As autophagy has been shown to play a key role in embryonic development and haematopoiesis ^30-32^ this provides further support to the CD44^Low^Kit^Neg^ cells being in transit from endothelial to haematopoietic cells.

### Runx1 is not required for the formation of CD44^Low^Kit^Neg^ endothelial cells

CD44 has allowed us for the first time to clearly define the key VE-Cad^+^ populations in the AGM. The transcription factor RUNX1 is a key driver of HSPC development and is known to down-regulate endothelial identity through its target genes GFI1 and GFI1B ^10,33^. Next, we decided to evaluate the impact of Runx1 loss of function on the different CD44^+^ cells. Using a Runx1 knockout mouse model ^34^, we stained for VE-Cad, CD44 and Kit expression and performed transcriptional profiling on the sorted cells (Fig. 7). We found that in the absence of Runx1 there is a loss of CD44^High^ and CD44^Low^Kit^Pos^ cells and we observed a concomitant increase in the frequency of the CD44^Low^Kit^Neg^ population (Fig. 7a and 7b). Interestingly, we found no obvious transcriptional differences between the CD44^Low^Kit^Neg^ populations derived from Runx1^+/+^ versus Runx1^-/-^ embryos indicating that Runx1 is not necessary for the formation of these endothelial cells but for the promotion of the transition into CD44^Low^Kit^High^ cells (Supplementary Fig. S5).

**Figure 7:**
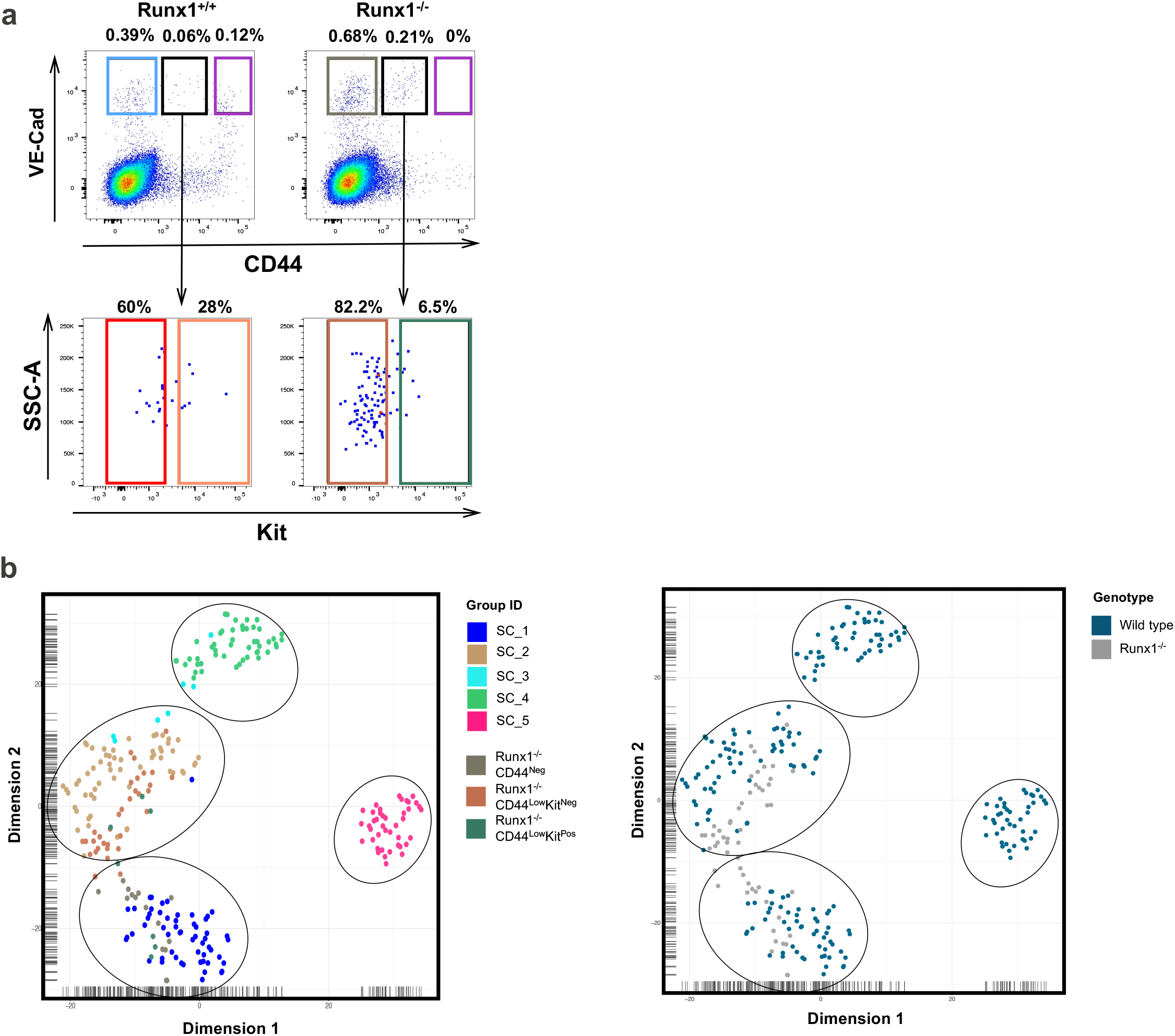
Runx1 is required for the generation of CD44^Low^Kit^Pos^ and CD44^Hlgh^ cells but not for the formation of the CD44^Low^Kit^Neg^ population. (**a**) FACS plots of VE-Cad and CD44 expression in the AGM region at E10.5 from Runx1^+/+^ (left) and Runx1^-/-^ (right) embryos. (**b**) tSNE plots from single-cell q-RT-PCR data shown in (a). See also Supplementary Fig. S5.

### All CD44^+^ populations display haematopoietic potential

To understand the haematopoietic potential of the different populations defined by VE-Cad, CD44 and Kit expression, we performed *ex vivo* assays using an OP9 co-culture system. No colonies were formed at the single-cell level from either the CD44^Low^Kit^Neg^ or the CD44^Neg^ populations. However, by plating cells at a density of 300 cells per well, we could observe round cell colony formation from the CD44^Low^Kit^Neg^ population but not from the CD44^Neg^ (Fig. 8a).

**Figure 8:**
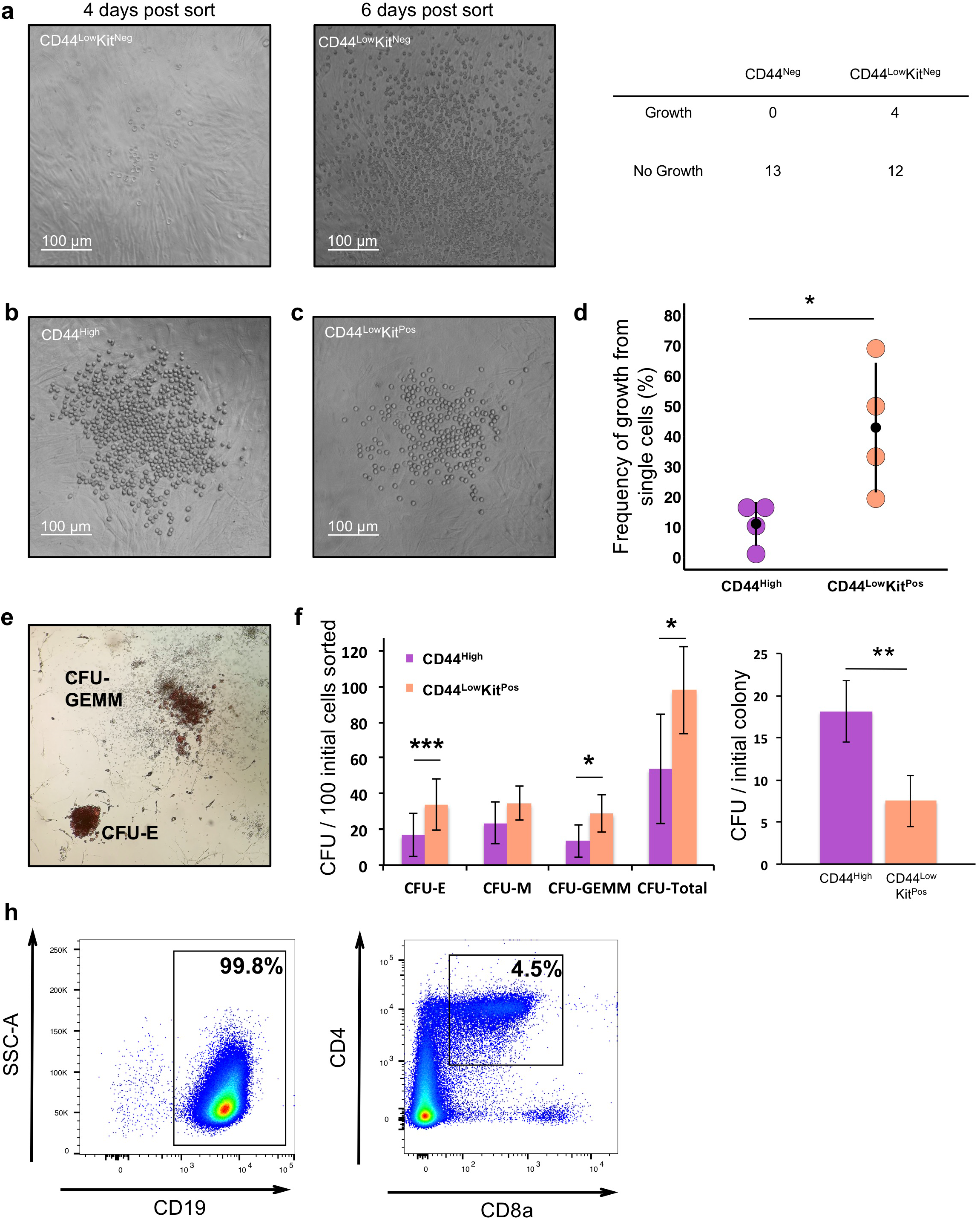
All CD44^+^ populations have haematopoietic potential. **(a)** Images of OP9 co-cultures after 4 and 6 days of incubation. Haematopoietic potential was observed from CD44^Low^Kit^Neg^ cells with colonies of round cells resulting from 300 cells sorted per well. No round cell colonies were observed with CD44^Neg^ cells. **(b)** Images of OP9 co-culture after 3 days of culture are shown. A single CD44^High^ or CD44^Low^Kit^Pos^ cell was FACS sorted onto a confluent OP9 stromal layer and incubated in HE medium. **(c)** The percentage of single cells giving rise to colonies was quantified across four independent experiment. The graph compares the frequency of growth from single CD44^High^ cells with CD44^Low^Kit^Pos^ cells. Statistical significance was determined by a two-tailed, paired t-test. **(d)** Colony forming unit assays were performed following three days OP9 culture of CD44^Low^Kit^Pos^ and CD44^High^ cells (100 cells per well). Cells were kept for a further 7 days in methocult medium before quantification. Images show representative CFU-E and CFU-GEMM colonies. **(e)** The bar graphs show the number of CFUs generated per 100 initial FACS sorted cells. Error bars indicate standard deviation (n=4). Significance was determined by two-tailed, paired t-tests. **(f)** The bar graph indicates the total number of colony forming units formed per initial colony grown on the OP9 stromal layer. Although the CD44^Low^Kit^Pos^ population gives rise to approximately 6 times more round cell colonies on OP9 then the CD44^High^ population, only approximately 2.5 times more CFUs are generated. **(g)** B and T cell lymphoid assays were performed following 21 days of OP9 (B-Cells) and OP9-DL1 (T-cells) culture of 50 FACS sorted CD44^High^ cells. Percentages of CD19^+^ (B-cells) and CD4^+^CD8a^+^ (immature T-cells) are shown.

In contrast, using single-cell sorting, we found that the CD44^Low^Kit^Pos^ population was the most potent with an average of 43% of single cells forming round cell colonies after three days of growth. Similarly, the CD44^High^ population produced haematopoietic colonies but with a lesser frequency, on average 11% of single cells showed the ability to form haematopoietic colony on OP9 (Fig. 8b-c). To uncover the differentiation potential of the cells generated by CD44^Low^Kit^Pos^ and CD44^High^, we performed CFU assays following the OP9 co-culture. Both populations readily generated both erythroid and myeloid colonies with the CD44^Low^Kit^Pos^ population demonstrating a significantly higher capacity than the CD44^High^ population (Fig. 8d-e). However, the four-fold increase in the number of CD44^Low^Kit^Pos^ colonies on OP9 did not correspond to a four-fold increase in CFU colonies suggesting a higher replating efficiency of the CD44^High^ cells compared to CD44^Low^Kit^Pos^ (Fig 8f). We further tested the lymphoid potential of the CD44^High^ population by growing cells for 21 days on either OP9 or OP9-DL1 with lymphoid promoting cytokines. We demonstrated that this population could give rise to both B and T cells *ex vivo* (Fig. 8h).

Overall, we found that all populations expressing CD44 displayed haematopoietic differentiation capacity including CD44^Low^Kit^Neg^ reinforcing the differentiation link between them as suggested by the transcriptome analyses described previously.

### Interrupting hyaluronan binding to the CD44 receptor inhibits the endothelial to haematopoietic transition *ex vivo* and *in vitro*

So far, we demonstrated that CD44 was a very useful marker to distinguish the different stages of EHT. Although the CD44 knock-out mice do not have a severe haematopoietic phenotype ^35^, compensatory mechanisms through other Hyaluronan receptors (e.g. Hmmr ^36^ expressed by some CD44^Low^Kit^Pos^ and CD44^High^ cells in Fig. 3a) may be at play to diminish the consequences of CD44 loss of function. In order to explore the functional role of CD44 in EHT we employed a pharmacological approach. By treating CD44^High^ sorted cells with a CD44 blocking antibody, we found that round cell colony formation could be inhibited in a dose dependent manner (Fig. 9a-b). The blocking antibody inhibited not only the number of colonies deriving from the *ex vivo* sorted cells but also the size of the colonies generated (Fig 9c).

**Figure 9:**
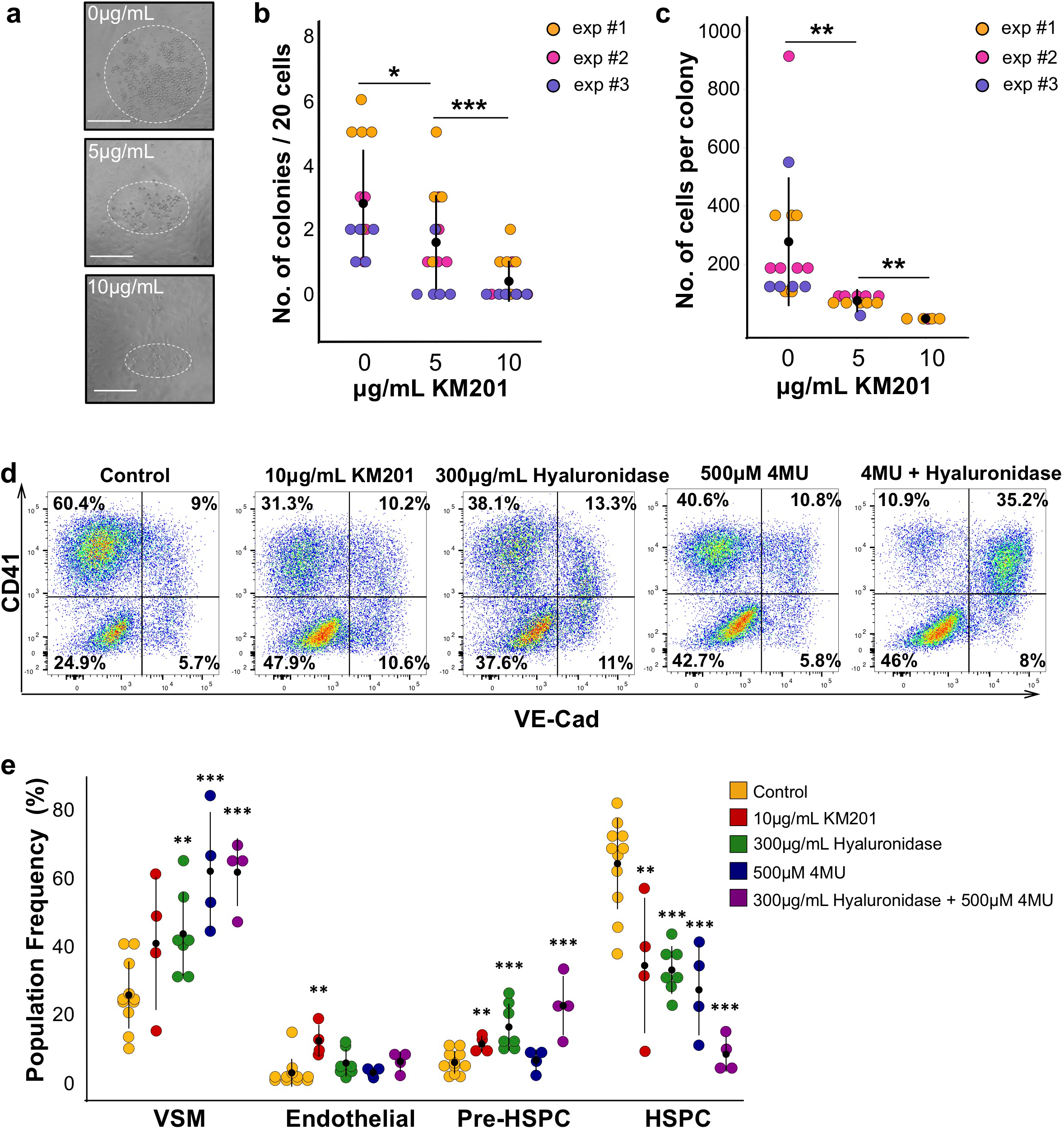
Blocking the interaction between CD44 and its ligand hyaluronan inhibits the endothelial-haematopoietic transition. (**a**) Images of round cell colonies generated from CD44^High^ cells after 4 days of OP9 co-culture with different concentrations of KM201 anti-CD44 blocking antibody. Dotted line indicates approximate size of colonies. Scale bar corresponds to 100 μm. **(b)** Dot plot comparing number of round cells colonies formed as a function of the concentration of anti-CD44 blocking antibody applied. **(c)** Dot plot indicating the number of cells per colony as a function of the concentration of anti-CD44 blocking antibody applied. **(b-c)** Significance was determined by two-tailed, independent t-test where * indicates p-value < 0.05, ** indicates p-value < 0.01 and *** indicates p-value < 0.001, n = 3. **(d)** Representative FACS plots of VE-Cad and CD41 expression after two days of haemangioblast differentiation. Cells were either untreated (control) or treated with anti-CD44 blocking antibody, hyaluronidase enzyme, 4MU hyaluronan synthase inhibitor or a combination **(e)** Dot plot showing the population percentage for vascular smooth muscle (VSM)(VE-Cad^-^CD41^-^), endothelial cells (VE-Cad^+^CD41^-^), Pre-HSPCs (VE-Cad^+^CD41^+^) and HSPCs (VE-Cad^-^CD41^+^) after two days of haemangioblast differentiation, summarising the results of FACS analysis shown in (d) (n ≥ 4). Significance was determined by two-tailed, independent t-tests where * indicates p-value < 0.05, ** indicates p-value < 0.01 and *** indicates p-value < 0.001. See also Supplementary Fig. S6.

To further investigate the role of CD44 in EHT we used an ESC *in vitro* differentiation system that mimics embryonic haematopoiesis. We performed a haemangioblast culture and analysed CD44 expression at day 1, 2 and 3. Endothelial cells expressed CD44 at all tested time points (Supplementary Fig. S6). We therefore applied the blocking antibody from the beginning of the culture and found that treatment with the antibody halved the number of HSPCs (VE-Cad^-^ CD41^+^) produced and significantly increased the percentage of endothelial cells in the culture (Fig. 8d-e). Given that the inhibitory antibody is known to bind close to the hyaluronan binding site on the extracellular domain of CD44 ^29^, we next attempted to manipulate the amount of hyaluronan in the culture. When 300 μg/mL of hyaluronidase was applied to the haemangioblast culture, we again observed a block in EHT characterised by a significant decrease in blood cell formation and an increase in the percentage of Pre-HSPCs (VE-Cad^+^ CD41^+^) (Fig. 8f). Using the 4MU inhibitor which blocks the synthesis of hyaluronan, we also observed the reduction of CD41^+^ cell number. Combining 4MU with hyaluronidase had a much more potent effect with a clear block in EHT at the Pre-HSPC stage.

In conclusion, these results for the first time demonstrate a regulatory role for hyaluronan and its receptor CD44 in the formation of HSPCs.

## Discussion

Using antibody screening and sc-RNA-seq to dissect the endothelial populations in the AGM, we discovered that CD44 was a robust marker to distinguish all the main populations in the EHT process. Combining CD44 with Kit and VE-Cad allowed us to discriminate the different stages of EHT more accurately that the method based on VE-Cad, CD41, CD43 and CD45 cell surface markers ^13^. In addition, we showed that CD44 had an unexpected regulatory function in the EHT process.

Our work has been instrumental in distinguishing the different types of endothelial cells in the AGM region. The CD44^Low^Kit^Neg^ has a gene expression signature strongly compatible with arterial identity (e.g. expression of *Efbn2* and *Sox17* and upregulation of Notch pathway target *Hey2)* while the CD44^Neg^ cells coexpressed genes related to the venous (e.g. *Nr2f2* and *Aplnr)* and arterial cell fates (e.g. *Efnb2* and *Sox17)* (Fig. 5d). This co-expression of venous and arterial genes supports previous work indicating that the dorsal aorta can contribute to the formation of the cardinal vein ^37^.

Another interesting finding was the striking metabolic state difference between the two endothelial populations. The CD44^Neg^ cells appeared much more metabolically active than the CD44^Low^Kit^Neg^ suggesting that the latter population is becoming quiescent. It is surprising since the acquisition of a quiescent phenotype in endothelial cells occurs normally after birth following the completion of angiogenesis. FOXO1 has been described as an important transcription factor to induce quiescence in endothelial cells through suppression of Myc ^38^. While we observe specific down-regulation of Myc in CD44^Low^Kit^Neg^ cells, *Foxo1* has a similar level of expression in the two endothelial populations. Recently, a study comparing lung endothelial cells between infant to adult mice showed that SMAD6 and SMAD7 higher expression in adult endothelial cells was linked with the induction of endothelial cell quiescence in adulthood ^39^. Interestingly, all the CD44^Low^Kit^Neg^ cells co-expressed *Smad6* and *Smad7* at the single-cell level suggesting that the quiescence phenotype we observed could also be linked to the co-expression of the two inhibitory Smads.

The acquisition of quiescence in this context could be the first indication of the divergence from an endothelial phenotype. The metabolic distinction of the CD44^Low^Kit^Neg^ population was coupled with an increase in genes involved in autophagy. This is in line with the known role of autophagy in embryogenesis, haematopoiesis and stem cell maintenance ^30-32,40,41^, such as protein and organelle turnover and protection from reactive oxygen species. Our data thus suggest that these cells mark the resetting of metabolic and regulatory states to initiate the EHT process.

We consequently propose that the CD44^Low^Kit^Neg^ population is the source of HSPCs in the AGM, hence the haemogenic endothelium (Fig. 10). Although it was suggested that HE cells in the human pluripotent stem cells differentiation model do not display arterial identity ^42^, our results clearly support the arterial source of HSPCs in the AGM. Another characteristic for the initiation of the EHT is the inhibition of the BMP and TGFbeta pathways as suggested by the increased expression of *Smad6, Smad7* and *Bmper* in the HE population. Expression of RUNX1 in endothelial cells would then trigger the haematopoietic cell fate. The dynamic interaction between the heptad of transcription factors GATA2, RUNX1, LYL1, ERG, FLI1, LMO2 and TAL1 co-expressed at the Pre-HSPC-I stage would eventually lead to the loss of endothelial gene expression ^24^ and give rise to the Pre-HSPC-II stage, cells expressing only haematopoietic genes.

**Figure 10:**
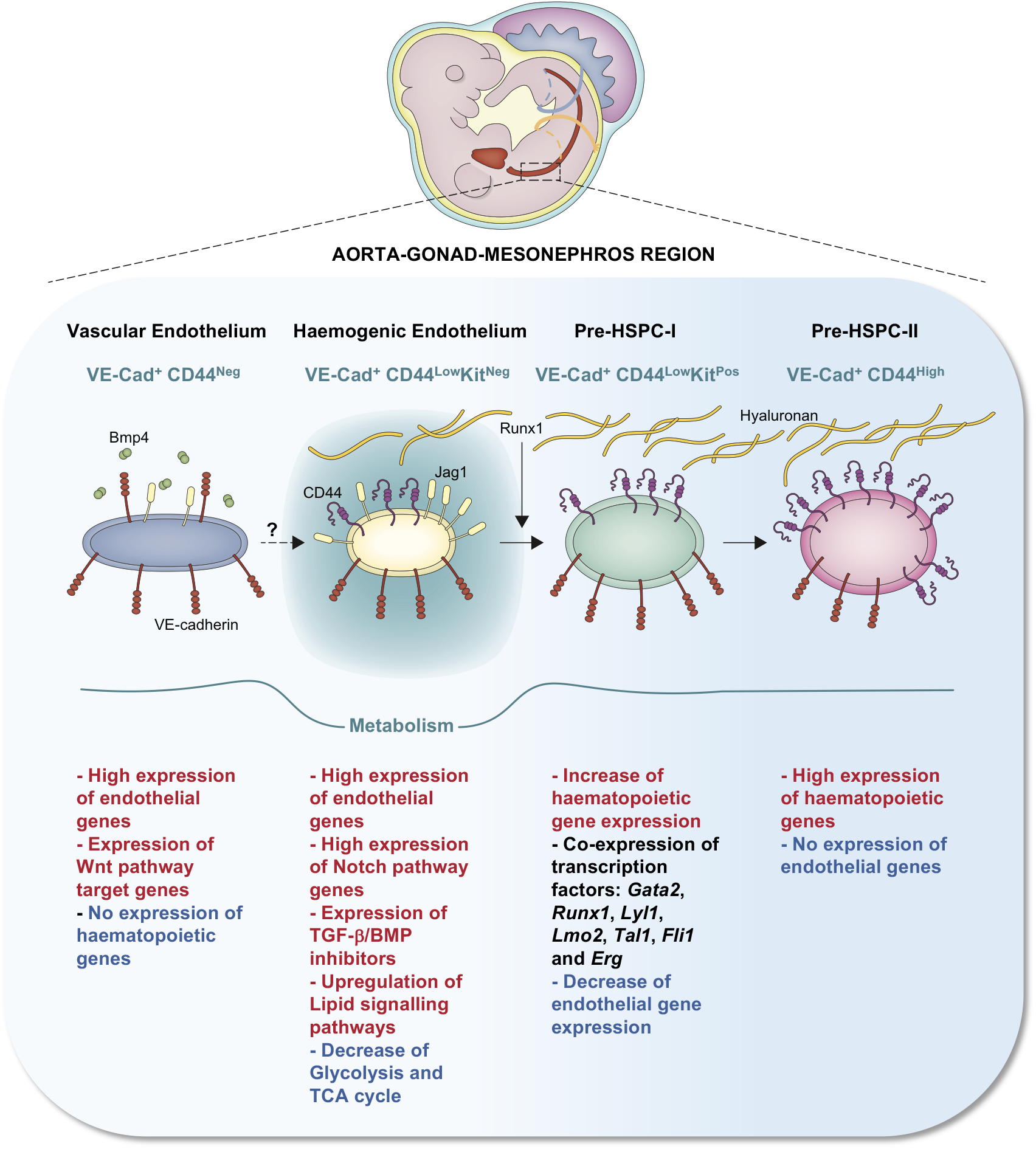
New model for the progression of EHT. Scheme summarising the findings of the present study. We identified two different sets of endothelial cell populations in the AGM with very distinct properties in term of signalling pathways and metabolic states. Expression of Runx1 in the haemogenic endothelium population triggers the upregulation of haematopoietic genes and the formation of the Pre-HSPC-I which co-expresses endothelial and haematopoietic genes. Continuous expression of haematopoietic genes and interaction between CD44 and hyaluronan eventually lead to the loss of endothelial genes and the formation of Pre-HSPC-II expressing only haematopoietic genes.

Our detailed transcriptomics analysis of hemogenic and non-hemogenic endothelial cell populations in the AGM revealed for the first time the prerequisites needed by endothelial cells to generate HSPCs. Thus, our study may help to design new protocols for the generation of HSCs *ex vivo* without the use of transcription factors, which still represents a considerable health risk. The manipulation of the interaction between CD44 and hyaluronan could offer a new strategy for reprogramming endothelial cells into HSPCs.

In addition, our work could shed new light on CD44 function in other cellular transitions such as the epithelial-mesenchymal transition occurring during metastasis ^23^. Given the role of CD44 in the metastatic process, it is possible that there is an overlap in its function for these transformations. Therefore, understanding the downstream targets of the CD44-hyaluronan interaction could also have implications also for cancer biology.

## Methods

### Timed mating and embryo dissection

For timed pregnancies, C57BL/6 wild-type mice or Runx1^+/-^ mice were mated overnight. Embryos were collected in PBS supplemented with 10% FBS (PAA Laboratories). The yolk sac of embryos derived from Runx1^+/-^ mice were genotyped using the Kappa Mouse Genotyping Kit (KAPA Biosystems) according to the manufacturer’s instructions. E9.5 to E11.5 embryos were staged based on somite counts. After removing the yolk sac, the AGM region was dissected by removing the head and tail above and below the limb buds then removing the limb buds, organs and somites by cutting ventrally and dorsally either side of the AGM.

All experiments were performed following the guidelines and regulations defined by the European and Italian legislations (Directive 2010/63/EU and DLGS 26/2014, respectively). They apply to foetal forms of mammals as from the last third of their normal development (from day 14 of gestation in the mouse). They do not cover experiments done with day 12 mouse embryos and at earlier stages. Therefore, no experimental protocol or license was necessary for the performed experiments. Mice were bred and maintained at the EMBL Rome Animal Facility in accordance with European and Italian legislations (EU Directive 634/2010 and DLGS 26/2014, respectively).

### C1 Fluidigm chip and single-cell RNA sequencing

AGM from E10 embryos were isolated and stained with anti-VE-Cadherin and anti-CD41 antibodies. VE-Cad^+^ cells from E10 embryos were isolated using FACS and mixed with Fluidigm suspension reagent (Fluidigm) in a 3:2 ratio. A primed Fluidigm C1 chip and 5ul of cell suspension was loaded onto a C1 instrument. Cell capture was then assessed with a 40x bright field microscope and wells scored as single-cell, doublet or debris. Lysis, reverse transcription and PCR reagents (Clontech Takara) along with ERCC spike-ins at 1 in 4000 dilution (Ambion) were added and mRNA-Seq RT & Amp script were performed overnight. The cDNA was then harvested and diluted in Fluidigm C1 DNA dilution buffer.

### Single-cell RNA sequencing data analysis

The paired 2 × 101bp Illumina reads from the libraries were quantified using Salmon ^43^ with the setting −l IU to indicate library topology, and the optional flags -- posBias and --gcBias to account for coverage and amplification biases present in sc-RNA-seq protocols. As an index the cDNA annotation of Ensembl release 85 for GRCm38.p4 was used, together with ERCC spike-in sequences. The TPM values were rescaled to not include ERCC expression and only consider endogenous gene expression.

Technical features of the data were compared with manual annotation of samples in C1 chambers through microscopy. Samples with less than 500,000 mapped reads and more than 30% mitochondrial content were discarded from analysis. This left 78 single cells in the *in vivo* experiment.

To identify cells as endothelial or haematopoietic, expression levels of 10 known markers were analysed (*Cdh5*, *Kdr*, *Pecam1*, *Pcdh12*, *Sox7*, *Gfi1*, *Gfi1b*, *Myb*, *Runx1*, *Spi1*). Cells were clustered using Principal Component Analysis and a Gaussian Mixture Model with two components on these markers (Fig. 1b). A cluster of 10 cells considered haematopoietic was identified (and annotated based on high Runx1 expression). Analysis was performed using the decomposition.PCA and mixture.GaussianMixture classes in the scikit-learn package. PCA was performed on the log transformed TPM values of the markers, and the first two principal components were used for Gaussian Mixture.

Novel markers for hematopoietic cells were identified using a likelihood ratio test, where the alternative model included a binary term for whether the cell was haematopoietic, and the null model just assumed a common mean for all the cells. The P-values from the likelihood ratio test were corrected for multiple testing by the Bonferroni procedure. The top differentially expressed genes were investigated to find markers which could be used in follow-up experiments, and *Cd44* was considered a good candidate (Fig. 1c).

### *In vitro* ES cell differentiation system

The A2lox Empty Embryonic Stem (ES) cell line was maintained and differentiated as previously described ^24^. Briefly, the ES cells were maintained on mouse embryonic fibroblasts in DMEM knockout medium (Gibco) supplemented with 1% Pen/Strep (Gibco), 1% L-glutamine (Gibco), 1% NEAA, 15% FBS (Gibco), LIF (EMBL Protein expression and purification core facility) and 2-Mercaptoethanol (Gibco).

For differentiation the cells were plated on 0.1% gelatin for 24 hours in DMEM-ES cell medium and for 24 hours with IMDM instead on DMEM-knockout and in the absence of NEAA. To generate embryoid bodies (EBs) ES cells were transferred to petri dishes at a density of 0.3×10^6^ cells per 10cm^2^ dish and plated in IMDM (Lonza) medium supplemented with 1% Pen/Strep, 1% L-glutamine (Gibco), 15% FBS, transferrin (Roche), MTG (Sigma) and 50mg/μL of ascorbic acid (Sigma). After 3 days in culture Flk1^+^ progenitor cells were isolated through magnetic activated cell sorting (MACS) using an anti-Flk1 APC conjugated antibody (eBiosciences) and anti-APC microbeads (Miltenyi Biotec).

For the haemangioblast differentiation, Flk1^+^ cells were cultured on gelatin for 24 to 72 hours in IMDM medium supplemented with 1% Pen/Strep, 1% L-glutamine (Gibco), 15% FBS (Gibco), transferrin (Roche), MTG (Sigma) and 50mg/μL of ascorbic acid (Sigma), 15% D4T, 10μg/mL VEGF (R&D), 10μg/mL IL6 (R&D). The cells could then be harvested with TrypLE express and cell populations analysed by flow cytometry using anti-VE-Cad, anti-CD41 and anti-Kit antibodies (supplementary table 1). For *in vitro* CD44-hyaluronan interaction experiments Flk1^+^ cells were plated on gelatin, as per the haemangioblast differentiation protocol, and incubated with either 10μg/mL anti-CD44 antibody [KM201], 300μg/mL of hyaluronidase (Sigma H4272), 500μM of 4MU (sigma). Generation of haematopoietic and endothelial populations was assessed by flow cytometry after 48hrs in culture.

### Antibody screen, Flow cytometry and cell sorting

The antibody screen was performed using the Mouse Cell Surface Marker Screening Panel (BD Bioscience, see table below) according to the manufacturer’s instructions. Cells from haemangioblast culture were harvested with TrypLE (Gibco) at 37 degrees for 5 minutes and deactivated with PBS supplemented with 10% FBS. Dissected AGMs were dissociated with collagenase (Sigma) at 37 degrees for 30 minutes and the collagenase deactivated with PBS supplemented with 10% FBS (Gibco). Following the generation of single cell suspension, cells were stained with different combination of antibodies (see list below). Cells were washed, filtered and analysed using FACS Aria III (Becton Dickinson) and FACS Diva software. Data was later analysed using FlowJo v10.1r5 (Tree Star Inc.). For single-cell qPCR analysis cells were sorted using an 85μm nozzle directly into reaction buffer (Cells Direct qRT-PCR Kit, Invitrogen) and snap frozen. For OP9 co-culturing assays cells were sorted using 100 μm nozzle directly into a 96 well culture dish (Costar).

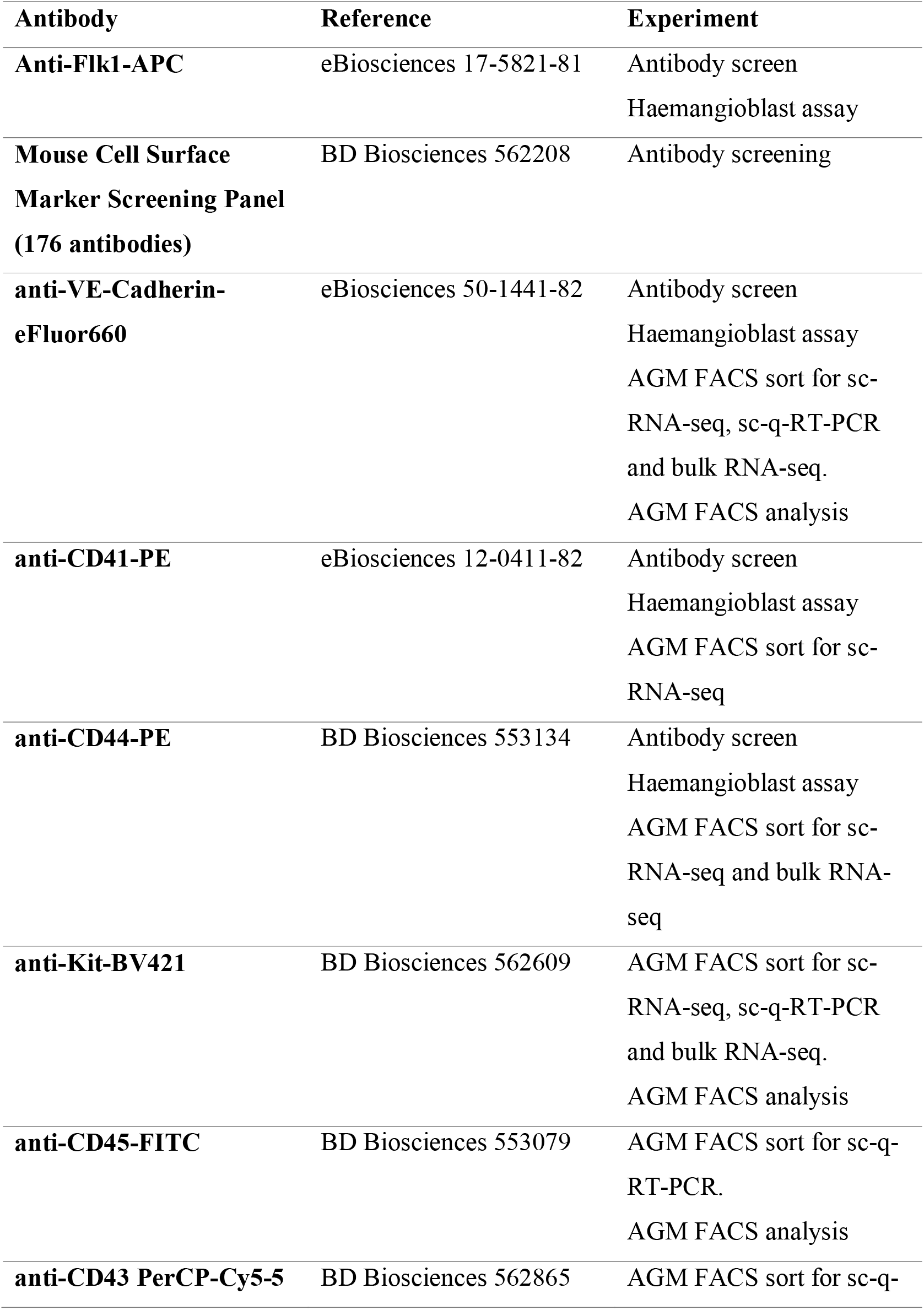
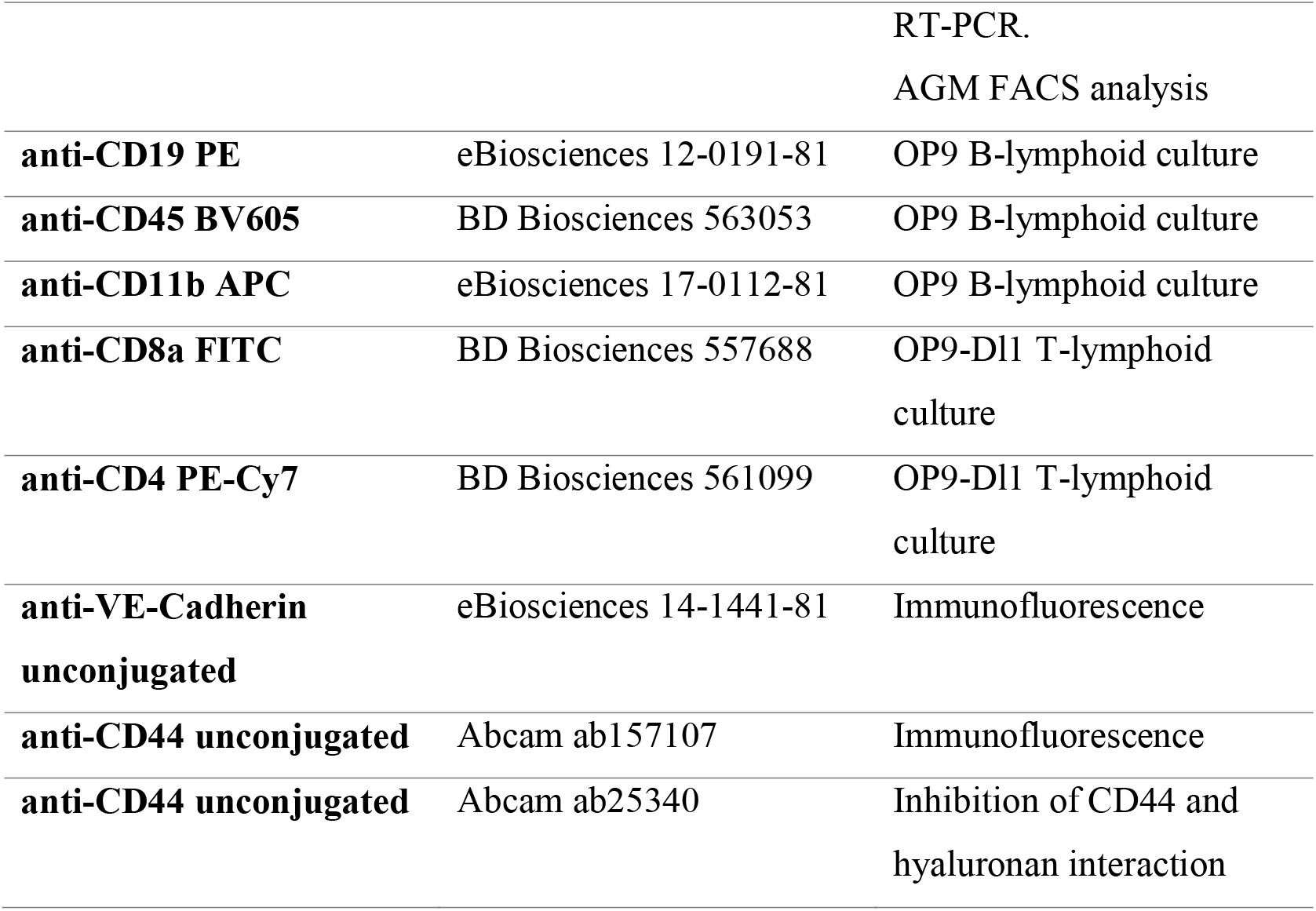

### Immunofluorescence and confocal microscopy

Mid-gestation embryos were dissected and fixed in 4% paraformaldehyde for 15 minutes at room temperature then incubated in 15% sucrose solution for 1 hour before freezing in OCT (Tissue-Tek). 10μm transverse cryo-sections of the AGM region were then placed on superfrost plus slides (Thermoscientific). Sections were washed in PBS, incubated in 1M glycine solution and permeabilised with 0.3% Triton X-100 (Sigma). Blocking solution consisting of 5% donkey serum, 5% chicken serum and 0.1% Tween-20 (Sigma) in TBS buffer was applied to sections for 2 hours at room temperature. Sections were incubated in primary antibodies overnight at 4°C and then washed. Secondary antibodies were applied for 1 hour at room temperature and washed before DAPI nuclear stain (Invitrogen) was applied for 15 minutes. Slides were washed before being mounted with Prolong gold (Life Technologies) and imaged on a Leica SP5 confocal microscope.

### Single-cell q-RT-PCR

Single cells were sorted directly into lysis buffer and snap frozen. Samples were reverse transcribed with superscript III reverse transcriptase from the Cells Direct one-step qRT-PCR Kit (Invitrogen) for 15 minutes at 50°C. The cDNA was then pre-amplified for twenty cycles with 25nM final concentration of each outer-primer for a set of 96 target genes (Supplementary Table S2). The cDNA was then diluted with loading reagent (Fluidigm) and SoFastTM EvaGreen supermix (Biorad) and loaded onto a chip with (50μM) of inner primer mix. Amplification of the target genes was measured with the Fluidigm Biomark HD system with the Biomark Data Collection software and the GE96 × 96 + Meltv2.pcl program.

### Single-cell qPCR data analysis

Analysis of single-cell qPCR data was performed as previously described ^24^. Briefly, initial analysis was performed using the Fluidigm Real Time PCR analysis software. Hierarchical clustering and principal component analysis was performed using the SINGuLAR analysis toolkit (Fluidigm version 3.5) in R software (version 3.2.1).

### Bulk RNA sequencing

Bulk RNA sequencing was performed as described by the SmartSeq2 protocol ^44^. Briefly, 25 cells from each group (see table below) were FACS sorted directly into lysis buffer containing 0.2% Triton X-100, oligo-dT primers and dNTP mix, and then snap frozen. Reverse transcription was then performed, followed by pre-amplification for 14 cycles. Nextera libraries were then prepared and sequenced on the Illumina Next Seq sequencer.

|  | **E9.5** | **E10** | **E11** |
| --- | --- | --- | --- |
| **VE-Cad<sup>Pos</sup> CD44<sup>Neg</sup>**<br>(CD44<sup>Neg</sup>) | 4 x 25-cells | 3 x 25-cells | 3 x 25-cells |
| **VE-Cad<sup>Pos</sup> CD44<sup>Low</sup> Kit<sup>Neg</sup>**<br>(CD44<sup>Low</sup> Kit<sup>Neg</sup>) | 1 x 25-cells | 2 x 25-cells | 3 x 25-cells |
| **VE-Cad<sup>Pos</sup> CD44<sup>Low</sup> Kit<sup>Neg</sup>**<br>(CD44<sup>Low</sup> Kit<sup>Neg</sup>) | 1 x 25-cells | 2 x 25-cells | Not done |
| **VE-Cad<sup>Neg</sup> CD44<sup>High</sup>**<br>(CD44<sup>High</sup>) | Not done | Not done | 3 x 25-cells |

### Bulk RNA-seq data analysis

Sequencing data was analysed with the aid of the EMBL Galaxy tools (galaxy.embl.de) ^45^ - specifically, FASTX for adaptor clipping, RNA STAR for mapping and htseq-count for obtaining raw gene expression counts. The R software (version 3.2.1, http://www.R-project.org) was then used to generate heatmaps and tSNE plots using the DESeq2, Scater, Biobase and pHeatmap packages.

### OP9 co-culturing assays

OP9 cells were maintained in MEM alpha medium (Gibco) with 20% FBS (ATCC 30-2020). For limiting dilution co-cultures cells were sorted directly onto a confluent OP9 stromal layer and incubated in medium conducive to haematopoietic development, IMDM (Lonza) treated with 5% penicillin-streptomycin (Gibco) and supplemented with 10% FBS (PAA Laboratories), L-glutamine, transferrin, MTG, ascorbic acid, LIF, 50ng/ml SCF, 25ng/ml IL3, 5ng/ml IL11, 10ng/ml IL6, 10ng/ml Oncostatin M, 1ng/ml bFGF. Round cell colonies were quantified after three to six days in culture. For *ex vivo* CD44 blocking antibody experiments 20 VE-Cad^+^CD44^High^ cells were sorted, as per OP9 co-culturing assays, into a medium containing either no antibody, 5μg/mL or 10μg/mL of anti-CD44 antibody [KM201] (Abcam). The number of colonies were assessed after 3 days in culture.

### Haematopoietic colony forming assay

One hundred cells were initially sorted onto a confluent OP9 stromal layer as per OP9 co-culturing assay. After three days in culture cells were harvested with TrypLE express (Gibco) and colony-forming unit-culture (CFU-C) assays were initiated using Methocult complete medium (Stem Cell Technologies). Cells were grown in 35mm culture dishes and colonies quantified after 7 days.

### Lymphocyte progenitor assay

Fifty cells were sorted onto confluent OP9 or OP9-DL1 stromal layers in MEM-alpha medium (Gibco) and supplemented with growth factors conducive to lymphocyte development, 20% FBS (PAA Laboratories), 50ng/ml SCF, 5ng/ml Flt-3L and 1ng/ml IL7. Medium was changed every 4-5 days and cells were split as necessary. Cells were cultured for 21 days before harvesting with TrypLE express for flow cytometry analysis.

## Author Contributions

Morgan Oatley, Conceptualization, Formal analysis, Investigation, Visualization, Writing—original draft, Writing—review and editing; Özge Vargel Bölükbas□i, Conceptualization, Formal analysis, Investigation, Visualization, Writing—review and editing; Valentine Svensson, Formal analysis, Visualization, Writing—review and editing; Maya Shvartsman, Investigation, Formal analysis, Writing—review and editing; Kerstin Ganter, Investigation, Visualization, Writing—review and editing; Katharina Zirngibl, Formal analysis, Visualization, Writing—review and editing; Polina V. Pavlovich, Formal analysis, Writing—review and editing; Vladislava Milchevskaya, Formal analysis, Writing—review and editing; Vladimir Foteva, Investigation, Writing—review and editing; Kedar N. Natarajan, Investigation, Writing—review and editing; Bianka Baying, Investigation, Writing—review and editing; Vladimir Benes, Supervision, Writing— review and editing; Kiran R. Patil, Supervision, Investigation, Writing— review and editing; Sarah A. Teichmann, Supervision, Investigation, Writing— review and editing; Christophe Lancrin, Conceptualization, Formal analysis, Supervision, Investigation, Visualization, Methodology, Writing—original draft, Project administration, Writing—review and editing.

## Acknowledgments

We thank Kalina Stantcheva and Cora Chadick (EMBL Rome FACS Facility, Italy) for cell sorting; Andreas Buness (EMBL Rome Bioinformatics Services, Italy) and Tallulah Andrews (Wellcome Trust Sanger Institute, UK) for help with bioinformatics analysis; Paul Collier, (EMBL Genomics Core Facility, Germany) for bulk RNA-seq; Gisela Luz (Patil Group, EMBL, Germany) for scientific illustration; Inke Nathke (University of Dundee, Scotland), Paul Heppenstall (EMBL, Italy) and Alexander Aulehla (EMBL, Germany) for fruitful scientific discussions. The European Molecular Biology Laboratory and the Wellcome Trust supported this work.

## References

1. Medvinsky, A. & Dzierzak, E. Definitive hematopoiesis is autonomously initiated by the AGM region. Cell 86, 897–906 (1996).

2. de Bruijn, M. F. T. R. et al. Hematopoietic stem cells localize to the endothelial cell layer in the midgestation mouse aorta. Immunity 16, 673–683 (2002).

3. Bertrand, J. Y. et al. Haematopoietic stem cells derive directly from aortic endothelium during development. Nature 464, 108–111 (2010).

4. Boisset, J.-C. et al. In vivo imaging of haematopoietic cells emerging from the mouse aortic endothelium. Nature 464, 116–120 (2010).

5. Kissa, K. & Herbomel, P. Blood stem cells emerge from aortic endothelium by a novel type of cell transition. Nature 464, 112–115 (2010).

6. Lancrin, C. et al. The haemangioblast generates haematopoietic cells through a haemogenic endothelium stage. Nature 457, 892–895 (2009).

7. Ciau-Uitz, A., Monteiro, R., Kirmizitas, A. & Patient, R. Developmental hematopoiesis: ontogeny, genetic programming and conservation. Experimental Hematology 42, 669–683 (2014).

8. Oberlin, E., Tavian, M., Blazsek, I. & Peault, B. Blood-forming potential of vascular endothelium in the human embryo. Development 129, 4147–4157 (2002).

9. Swiers, G. et al. Early dynamic fate changes in haemogenic endothelium characterized at the single-cell level. Nature Communications 4, 1–10 (2014).

10. Thambyrajah, R. et al. GFI1 proteins orchestrate the emergence of haematopoietic stem cells through recruitment of LSD1. Nat Cell Biol 18, 21–32 (2016).

11. Sugimura, R. et al. Haematopoietic stem and progenitor cells from human pluripotent stem cells. Nature 545, 432–438 (2017).

12. Lis, R. et al. Conversion of adult endothelium to immunocompetent haematopoietic stem cells. Nature 545, 439–445 (2017).

13. Rybtsov, S. et al. Tracing the origin of the HSC hierarchy reveals an SCF-dependent, IL-3-independent CD43(-) embryonic precursor. STEMCR 3, 489–501 (2014).

14. Zhou, F. et al. Tracing haematopoietic stem cell formation at single-cell resolution. Nature 533, 487–492 (2016).

15. Aruffo, A., Stamenkovic, I., Melnick, M., Underhill, C. B. & Seed, B. CD44 is the principal cell surface receptor for hyaluronate. Cell 61, 1303–1313 (1990).

16. Takaishi, S. et al. Identification of gastric cancer stem cells using the cell surface marker CD44. STEM CELLS 27, 1006–1020 (2009).

17. Prince, M. E. et al. Identification of a subpopulation of cells with cancer stem cell properties in head and neck squamous cell carcinoma. Proceedings of the National Academy of Sciences 104, 973–978 (2007).

18. Lopez, J. I. et al. CD44 attenuates metastatic invasion during breast cancer progression. Cancer Research 65, 6755–6763 (2005).

19. Liu, C. et al. The microRNA miR-34a inhibits prostate cancer stem cells and metastasis by directly repressing CD44. Nature Medicine 17, 211–215 (2011).

20. Du, L. et al. CD44 is of functional importance for colorectal cancer stem cells. Clin. Cancer Res. 14, 6751–6760 (2008).

21. Avigdor, A. et al. CD44 and hyaluronic acid cooperate with SDF-1 in the trafficking of human CD34+ stem/progenitor cells to bone marrow. Blood 103, 2981–2989 (2004).

22. Zovein, A. C. et al. Fate tracing reveals the endothelial origin of hematopoietic stem cells. Cell Stem Cell 3, 625–636 (2008).

23. Pastushenko, I. et al. Identification of the tumour transition states occurring during EMT. Nature 556, 463–468 (2018).

24. Bergiers, I. et al. Single-cell transcriptomics reveals a new dynamical function of transcription factors during embryonic hematopoiesis. eLife 7, R106 (2018).

25. Quintero, M., Colombo, S. L., Godfrey, A. & Moncada, S. Mitochondria as signaling organelles in the vascular endothelium. Proceedings of the National Academy of Sciences 103, 5379–5384 (2006).

26. Yu, X., Long, Y. C. & Shen, H.-M. Differential regulatory functions of three classes of phosphatidylinositol and phosphoinositide 3-kinases in autophagy. Autophagy 11, 1711–1728 (2015).

27. Nascimbeni, A. C., Codogno, P. & Morel, E. Phosphatidylinositol-3-phosphate in the regulation of autophagy membrane dynamics. FEBS J. 284, 1267–1278 (2017).

28. Tan, X., Thapa, N., Liao, Y., Choi, S. & Anderson, R. A. PtdIns(4,5)P2 signaling regulates ATG14 and autophagy. Proc. Natl. Acad. Sci. U.S.A. 113, 10896–10901 (2016).

29. Dall’Armi, C., Devereaux, K. A. & Di Paolo, G. The role of lipids in the control of autophagy. Curr. Biol. 23, R33–45 (2013).

30. Riffelmacher, T. & Simon, A. K. Mechanistic roles of autophagy in hematopoietic differentiation. FEBS J. 284, 1008–1020 (2017).

31. Phadwal, K., Watson, A. S. & Simon, A. K. Tightrope act: autophagy in stem cell renewal, differentiation, proliferation, and aging. Cell. Mol. Life Sci. 70, 89–103 (2013).

32. Mizushima, N. & Levine, B. Autophagy in mammalian development and differentiation. Nat Cell Biol 12, 823–830 (2010).

33. Lancrin, C. et al. GFI1 and GFI1B control the loss of endothelial identity of hemogenic endothelium during hematopoietic commitment. Blood 120, 314–322 (2012).

34. Pinheiro, I. et al. Discovery of a new path for red blood cell generation in the mouse embryo. bioRxiv 309054 (2018). doi:10.1101/309054

35. Schmits, R. et al. CD44 regulates hematopoietic progenitor distribution, granuloma formation, and tumorigenicity. Blood 90, 2217–2233 (1997).

36. Nedvetzki, S. et al. RHAMM, a receptor for hyaluronan-mediated motility, compensates for CD44 in inflamed CD44-knockout mice: A different interpretation of redundancy. Proceedings of the National Academy of Sciences 101, 18081–18086 (2004).

37. Lindskog, H. et al. Molecular identification of venous progenitors in the dorsal aorta reveals an aortic origin for the cardinal vein in mammals. Development 141, 1120–1128 (2014).

38. Wilhelm, K. et al. FOXO1 couples metabolic activity and growth state in the vascular endothelium. Nature 529, 216–220 (2016).

39. Schlereth, K. et al. The transcriptomic and epigenetic map of vascular quiescence in the continuous lung endothelium. eLife 7, 464 (2018).

40. Mortensen, M. et al. The autophagy protein Atg7 is essential for hematopoietic stem cell maintenance. Journal of Experimental Medicine 208, 455–467 (2011).

41. Ho, T. T. et al. Autophagy maintains the metabolism and function of young and old stem cells. Nature 543, 205–210 (2017).

42. Ditadi, A. et al. Human definitive haemogenic endothelium and arterial vascular endothelium represent distinct lineages. Nat Cell Biol 17, 580–591 (2015).

43. Patro, R., Duggal, G., Love, M. I., Irizarry, R. A. & Kingsford, C. Salmon provides fast and bias-aware quantification of transcript expression. Nature Methods 14, 417–419 (2017).

44. Picelli, S. et al. Full-length RNA-seq from single cells using Smart-seq2. Nature Protocols 9, 171–181 (2014).

45. Afgan, E. et al. The Galaxy platform for accessible, reproducible and collaborative biomedical analyses: 2016 update. Nucleic Acids Research 44, W3–W10 (2016).

